# Multi-omics dissection of Parkinson’s patients in subgroups associated with motor and cognitive severity

**DOI:** 10.1101/2025.02.20.638984

**Authors:** Efi Athieniti, Sotiroula Afxenti, George Minadakis, George M. Spyrou

**Author notes:** Contributing authors.

## Abstract

Heterogeneity in the severity of Parkinson’s disease (PD) inhibits the effective interpretation of clinical trial outcomes. Multi-omics analysis may help explain the pathological mechanisms underlying disease progression and reveal biomarkers of clinical severity. We performed Multi-Omics Factor Analysis (MOFA) on whole blood RNA, miRNA and Cerebrospinal fluid (CSF) and blood plasma proteomics from the Parkinson’s Progression Marker Initiative (PPMI), to identify molecular factors correlated with motor (MDS-UPDRS3) and cognitive (Semantic Fluency Test, SFT) function. Three molecular factors significantly correlated with the MDS-UPDRS3 score and two with SFT, which remained significant after adjusting for age, sex, and medication dose. We used the identified factors to stratify patients into subgroups with distinct motor and cognitive severity. The severe motor clusters showed deregulation of cytotoxic natural killer cell mechanisms in peripheral blood, and changes to proteins associated with the endoplasmic reticulum in CSF. The severe cognitive clusters showed changes in complement and synaptic dysfunction. Our analysis capitalizes on multi-omics data integration to enrich our understanding of the mechanisms driving motor and cognitive decline in PD, to support precision medicine.

## 1 Introduction

Parkinson’s disease (PD) is the 2^nd^ most common neurodegenerative disorder and its main pathological hallmark is the progressive loss of dopaminergic neurons in the substantia nigra of the brain [1]. This region is responsible for motor coordination and its degeneration leads to the major motor symptoms of PD, resting tremor, bradykinesia, rigidity and postural flex disturbance. In addition to these, nonmotor symptoms also develop, including cognitive decline, major neuropsychiatric symptoms, autonomous system disorders, sleep disorders, and sensory symptoms.

Idiopathic PD is multifaceted, involving various pathological mechanisms such as the misfolding of *α*-synuclein, the mitochondrial dysfunction, autophagy and oxidative stress [2]. This complexity makes both the diagnosis and the development of effective therapies difficult. Clinical trials have achieved limited success partly because of the lack of reliable markers for assessing disease severity. For example, interventions targeting the immune system have moved forward, but most trials focus on clinical rather than immunological endpoints. Integrating immune-related endpoints may be an important consideration to accurately assess the success of the trial [3]. There are major drawbacks to having a staging system based only on clinical measures with scales and questionnaires, such as subject-to-inter-rater variability, confounded by symptomatic drug effects, non-linear changes over time and more [4]. Therefore, there is growing emphasis on establishing a biological staging system of the disease based on molecular biomarkers.

There is active research in identifying biomarkers with diagnostic, prognostic, predictive, monitoring and pharmacodynamic value in PD. Recently, a study by the Parkinson’s Progression Markers Initiative (PPMI) has presented a new biomarker assay, namely the *α* -synuclein seed amplification assay (SAA) which shows promise in differentiating PD patients from Healthy controls (HC) [5]. However, there are still no biomarkers established to be used in monitoring the progression of the disease, which may be different from diagnostic biomarkers. Diagnostic biomarkers may not change during disease progression and, therefore, may be informative regarding the initiation of disease but not as useful for monitoring disease progression [6]. Monitoring biomarkers can be measured longitudinally to assess changes in the severity of the disease and the effectiveness of therapy, which would significantly improve the outcome of clinical trials.

The difficulty in assessing the progression of patients is also a factor that inhibits the accurate estimation of trial outcomes. Within PD there is large heterogeneity, and patients exhibit a significant variation in the progression rates of motor, non-motor and psychological symptoms. Identifying the molecular markers that explain these rates can help recruit the patients who would have the most benefit from the treatments. So far, many subtyping methods applied do not have established therapeutic implications. This may not be surprising given that they were born largely from clinical observations of phenotype and not in observations regarding treatment response or biological hypotheses [7].

Previous studies have shown that there might be a molecular signature associated with disease progression in peripheral fluids such as blood and plasma. Several inflammatory markers such as TNF-*α*, IL-6 and IL-1*β* are found to be elevated as the disease progresses, and associated with worse motor and cognitive function [8]. Reduced plasma levels of protein DJ-1, which protects neurons from oxidative stress, have been associated with disease progression [9]. Increased levels of Neurofilament Light Chain (NFL) protein, a marker of neuronal damage, but also a prominent component of abnormal intraneuronal aggregates have been linked to the severity and progression of PD [10].

Recently, multi-omics analysis has shown promise in understanding disease and predicting disease states. For instance, a multi-panel approach using various proteins showed improved performance and robustness across datasets compared to single protein analyses [11]. By integrating genomics and transcriptomics studies have fine-mapped PD-associated loci, revealing specific variants that impact gene regulation and disease susceptibility. In the example in [12], two novel SNPs were identified, rs7294619 which disrupts LRKK2 expression via FGCGR2A, and rs4771268 which affects the MBNL2 regulation enhancer in oligodendrocytes. Furthermore, multi-omics approaches can provide insights into cross-layer molecular mechanisms. For example, a review of the use of multi-omics datasets from PD highlighted that neuroinflammation is consistently observed across multiple omic layers, reinforcing the confidence in its role in disease pathology [13].

Clustering analyses have also been used to identify subtypes of Parkinson’s disease (PD) patients, but these have largely relied on clinical features, progression patterns, or single-modality molecular data. For example, [14] utilized similarity fusion clustering on BioFIND data, identifying subtypes with distinct motor and non-motor severity profiles but found no significant differences in molecular markers or genetics across subtypes. The study in [15] used clinical and biomarker data to explore progression patterns, identifying subtypes defined by distinct disease trajectories. Similar analysis of early-onset PD revealed clinical subtypes based on motor and non-motor features, emphasizing the heterogeneity in disease presentation [16]. While, these approaches highlight the value of clustering in understanding PD heterogeneity, they are limited by their reliance on clinical data or single omics layers.

To overcome these limitations, global analysis methods like multi-omics factor analysis have emerged as powerful tools for investigating disease mechanisms and patient heterogeneity. By focusing on the covariation of molecular modules, factor analysis can disentangle biological and technical variation in omics data. Moreover, it can prioritize molecular features with the highest variation associated with clinical phenotypes, enabling the identification of molecular subgroups with distinct disease characteristics. These methodologies have recently shown promise in uncovering novel disease subtypes and mechanisms in various diseases [17, 18]. Despite this potential, molecular multi-omics clustering has not yet been applied to PD to identify patient subgroups and connect them to clinical severity.

Recognizing the gap in multi-omics studies, we leveraged a multi-omics dataset from the Parkinson’s Progression Markers Initiative (PPMI) [19], to explore mechanisms related to the progression of the disease. We integrated blood RNA, miRNA and plasma and CSF proteomics data, to uncover novel PD subtypes and their underlying molecular mechanisms, providing a comprehensive perspective on the disease’s severity profile. Our hypothesis is that integrating the various omics datasets that have been studied independently so far, can provide a more comprehensive picture of the molecular mechanisms associated with disease progression that span through the various molecular layers and tissues.

In this paper, we present a computational study on the multi-omics dataset from the PPMI aiming to detect biological stages of the disease. To achieve this, the analysis has completed the following objectives:

- Apply multi-omics factor analysis to detect molecular factors correlating with the disease severity.
- Use these molecular factors to cluster samples and identify PD subgroups with varying severity.
- Characterise the variations in molecular markers across subgroups to gain insights into disease progression.
- Apply computational drug repurposing exploiting the molecular markers to recommend drugs

Our workflow overview is depicted in Fig. 1. Using molecular profiles collected at year 3 post-diagnosis, when disease severity states are well established, we applied Multi-Omics Factor Analysis (MOFA) [20] to identify molecular factors correlating with motor (MDS-UPDRS3) and cognitive (Semantic Fluency Test, SFT) severity. Patients were clustered into subgroups based on these factors, revealing distinct clinical severity patterns. Differential expression and pathway enrichment analyses identified key RNA molecules, proteins, and pathways linked to disease severity. Furthermore, we applied computational drug repurposing and highlighted potential therapeutic targets based on the molecular findings. This integrative, multi-modal approach provides insights into the molecular heterogeneity of PD and its relationship to clinical progression.

**Fig. 1:**
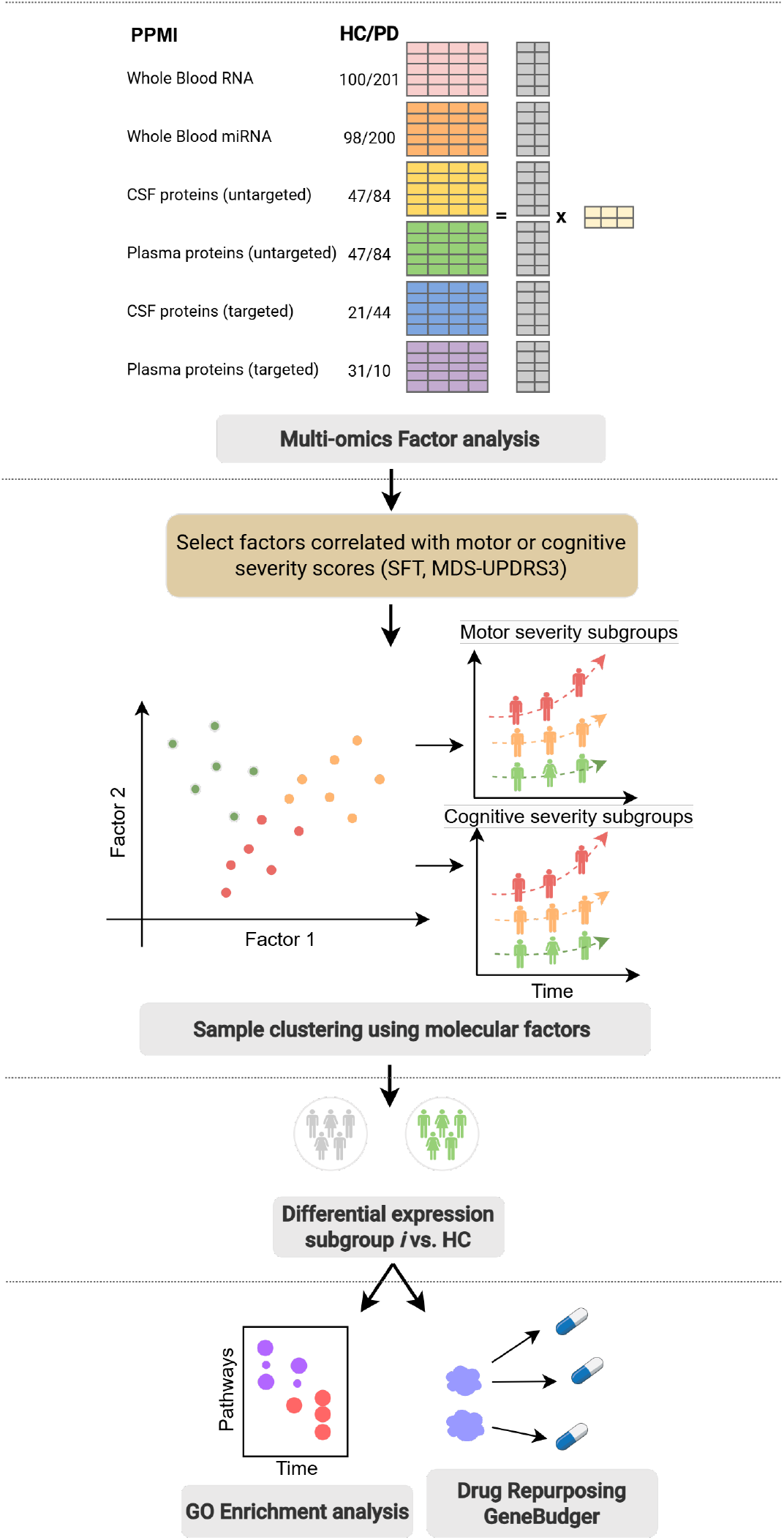
Overview of the methodology. Six omic datasets from year 3 were input to MOFA. The factors correlated with clinical scores were used to cluster PD samples. Each of the subgroups underwent differential expression analysis for each omics dataset to detect the top DE molecules between each subgroup and HC. The values across the first 3 years of study were obtained for the top DE markers per subgroup to understand temporal variations.

## 2 Results

### 2.1 Multi-omics Factor analysis results

The multi-factor analysis model captured 75%, 56%, 33%, 8%, 46%, and 49% of the variance from the RNA, miRNA, untargeted CSF proteomics, untargeted plasma proteomics, targeted CSF proteomics and targeted plasma proteomics. To detect molecular factors associated with clinical scores, the factor values of PD samples were tested for correlations with their clinical scores (Fig. 2). MDS-UPDRS3 significantly correlated with factors 13, 20 and 28, with *r* = 0.2, 0.19, − 0.19, respectively. The SFT score showed a correlation with factors 13, 16, 23 with values of *r* = − 0.32, 0.17 and 0.18. To check that the factors remained significant predictors of the scores after correcting for confounding variables, we performed linear regression analysis of each score and factor with age, sex and LEDD as covariates. All factors remained significant for MDS-UPDRS3, and for SFT only 13 and 16 remained significant after corrections (see Supplementary Fig. 1 for model parameters and fit).

**Fig. 2:**
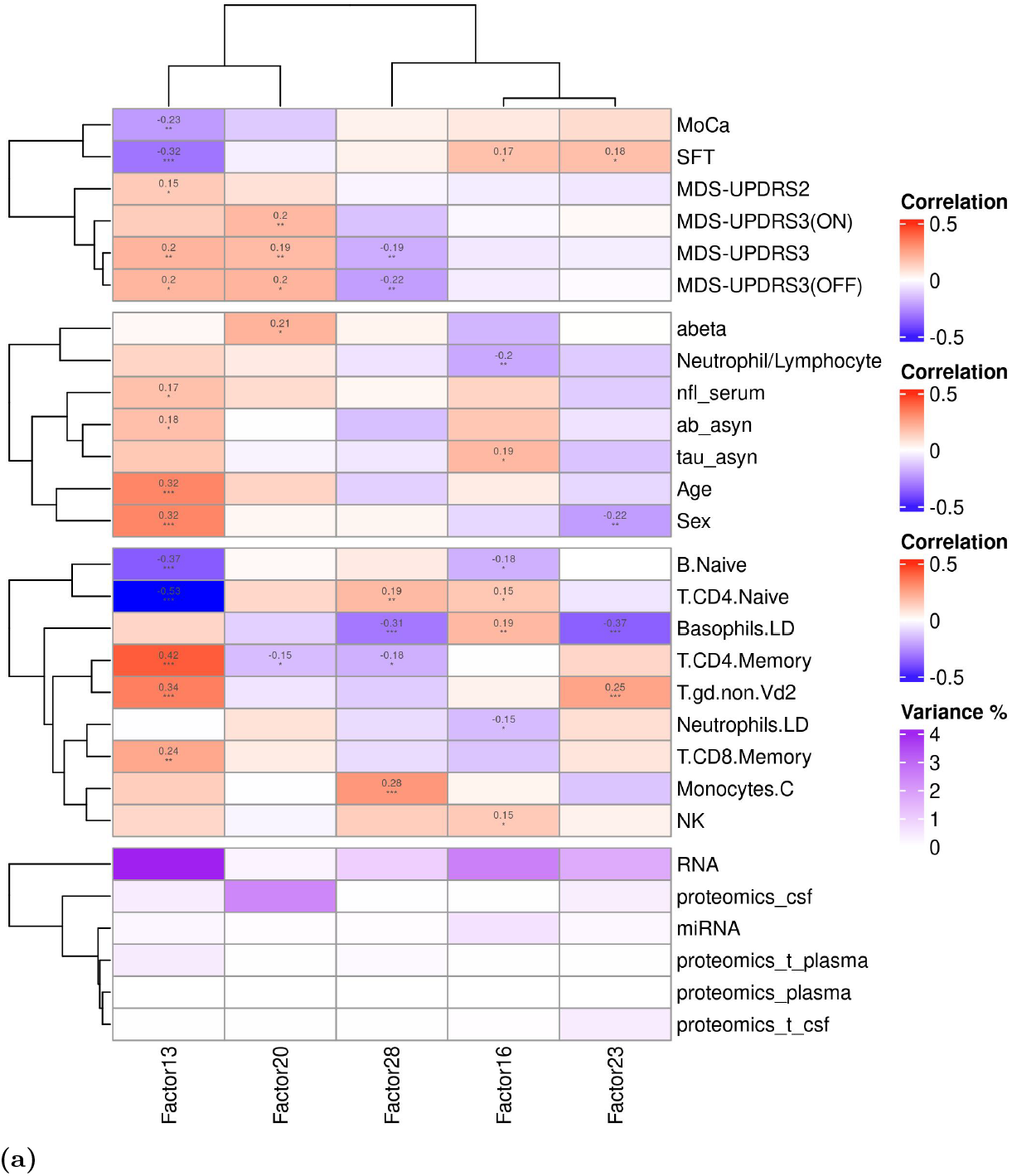
Factor annotations. Correlations of factors with selected clinical scores, clinical markers, covariates, and blood cell type proportions for PD patients. MOFA variance explained by factor and modality. Only correlations for variables that were significant (p<0.05) are shown.

#### Factor annotation: Clinical Scores, Blood cell type compositions and enriched pathways

To help understand the biological processes represented by the factors that showed significant correlations with the clinical scores MDS-UPDRS3 and SFT, we discuss here, the top two omics modalities contributing to the variance explained by the factor, the highly correlated clinical variables (Fig. 2), and the most highly enriched pathways (Supplementary Fig. 2) from the top modalities of each factor.

#### Factor 13: Age, Sex, Natural killer and B-cell immunity

This factor is associated with age and sex, ab asyn and both of the clinical scores tested MDS-UPDRS3 and SFT. The top modalities are RNA and targeted plasma proteomics. Among the molecular pathways enriched in the RNA modality are natural killer cell-mediated immunity (*KNKG7, KLRD1, GZMB*), cell killing (*NKG6, KLRD1, GNLY*) and B cell receptor signalling (*IGHM, CD22, BANK1*). Gamete generation (SRC, TARBP2, HPGDS), glutamatergic synapse (SRC, IL1RAP, PRKR1A) and regulation of small GTPase mediated signaling transduction (SRC, ARHGAP1, ADCYAP1R1) is enriched in targeted plasma proteomics.

#### Factor 20: Composition of CD4 memory cells, Lymphocyte activation and CSF protein localization

This factor correlated with CD4mem cell type composition, capturing variance primarily from untargeted CSF proteomics followed by blood RNA. The enrichment analysis from untargeted CSF proteomics showed protein localization to extracellular region (VGF, CHGA, FGG) and regulation of secretion (CHGA, FGG, FGA) and blood RNA was enriched with the structural constituent of ribosome (*RPL9, RPS18, RPS6*) and cytoplasmic translation (*RPL9, RPS18, RPS6*).

#### Factor 28: Composition of CD4 naive and memory cells, Leukocyte apoptosis and cell adhesion molecules

It is associated with the composition of CD4 naive (CD4nv), CD4 memory (CD4mem), Basophils and Monocytes. The top modalities are RNA and targeted plasma proteomics. It is enriched with activated T-cell proliferation (*CDC, IDO1, CD274*) and G-protein coupled receptor activity (*PTGDR2, HRH4, ADGRE1*) in blood RNA, SH3 domain binding (PTPN6, HCLS1, LYN) and leukocyte apoptotic process (HCLS1, MIF, LYN, FADD) in targeted plasma proteomics. The above shows that there are similar inflammatory processes detected both in blood and plasma.

#### Factor 16: CSF tau/*α*-synuclein, composition of B naive and CD4 naive cells and virus response

It is correlated with the measured Neutrophil to Lymphocyte ratio and it captures variance from RNA and miRNA. It has negative correlation with the composition of B-naive (Bnv) cells and neutrophils and positive with CD4nv, Basophils, NK cells. In the RNA modality, it is enriched with defense response to virus (*IFI44L, RSAD2, IFI44*) and platelet alpha granule (*ITGB3, CLU, CD9*), and in targeted CSF proteomics with positive regulation of protein catabolic process (SIRT2, HSPA1A, PTEN) and neuron projection guidance (NCAM1, CHL1, DAG1). There are no significantly enriched pathways from other modalities.

Linear regression analysis was performed for all blood cell type compositions and clinical scores that correlated with the factor, evaluating their direct association with the scores. Among the various covariates, only the composition of Bnv showed a significant negative association with the MDS-UPDRS3 score (Supplementary Fig. 1) Basophils and CD8 naive (CD8nv) cells decreased as the SFT score decreased, indicating a relationship with increased cognitive severity.

### 2.2 Clustering patients with molecular multi-omics data reveals subgroups with distinct clinical severity and progression patterns

#### 2.2.1 Clustering patients based on factors correlating with motor examination (MDS-UPDRS3)

##### Overview of clustering approach and clinical correlations

Three PD groups were constructed using k-means clustering of the multi-omics factors, resulting in n=97, 59 and 38 patients in each group respectively. The multi-omics factors effectively produced subgroups with significant differences in clinical scores. A qualitative heatmap of the medians of the clinical scores in each group is shown in Fig. 3, and a quantitative table is shown in Table 3. Cluster 1 and 2 have a higher median MDS-UPDRS3 at year 3 (Wilcoxon signed rank test, *p* < 0.05) compared to cluster 3 and HC. Based on the MDS-UPDRS3 score observed, the clusters were labelled as follows Cluster 1: Severe motor (SM), Cluster 2: Intermediate motor (IM), and Cluster 3: Mild motor (MM). SM and IM are predominantly male: 77% and 81%, vs. 54% of cluster MM. The average age is lower in cluster MM (60.15) and then SM (64) and IM (64.5). Cluster SM and IM also show trends of worse severity in other scales: lower MOCA and HVLT immediate recall and higher SCOPA. Cluster IM also shows the highest Geriatric Disease Scale (GDS) score.

**Fig. 3:**
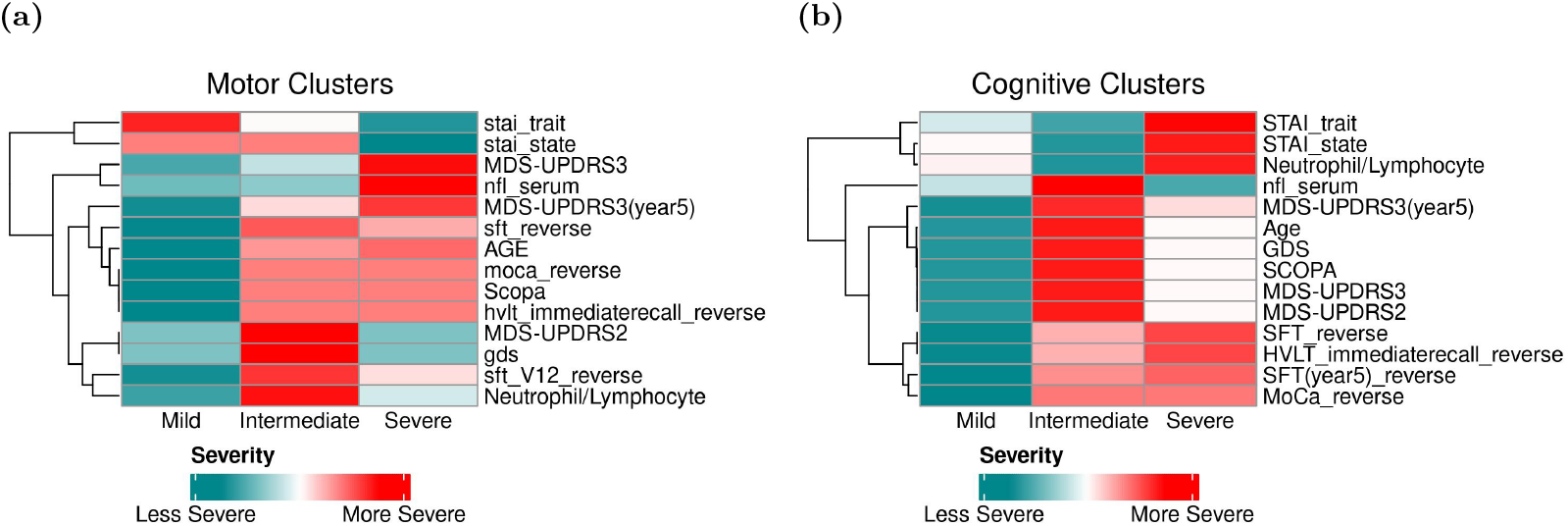
Qualitative summary of the clinical scores (z-score) for the three clusters obtained using the molecular factors associated with (a) MDS-UPDR3 and (b) SFT. The clusters are positioned by order of severity of the clinical score MDS-UPDRS3 and SFT.

##### Top differentially expressed molecular markers and pathways between Motor severity subgroups

To understand the molecular variations that led to the clusters produced, we first inspected the factor values for each cluster (Supplementary Fig. 4). The severe and intermediate motor clusters take higher values of factor 13 compared to the mild motor cluster. Cluster IM also takes a higher value in factor 20 and cluster SM takes a lower value of factor 28 compared to the others. This indicates that different molecular factors and thus biological processes were identified as contributing to the various clusters with differences in motor severity. The differential expression analysis (DEA) of each cluster compared to HC resulted in 251, 66, 692 (Supplementary Dataset 1) differentially expressed RNA molecules in cluster SM, IM and MM respectively. The proportions of blood cell types estimated from the RNA expression dataset (Supplementary Fig. 3, Supplementary Table 2), showed that the severe motor group had depleted CD4nv and Bnv cells compared to HC and MM. It is also interesting to note that MM has significantly lower CD4mem cells, both compared to the severe clusters and HC. We repeated DEA with the cell type compositions as covariates in the model, which dramatically reduced the number of differentially expressed molecules to 0, 1 and 4 in cluster SM, IM and MM respectively, showing that the largest variation in the RNA sequencing modality is contributed by variability in the composition of immune cell types.

##### RNA Differences relative to Healthy Controls

To obtain an overview of the differences in gene expression patterns we performed GSEA. The top enriched pathways compared to HC as obtained by ClusterProfiler are shown in Supplementary Fig. 6. They include cell killing, chemotaxis, secretory granule lumen, ribosome biogenesis, ncRNA processing, mitochondrial gene expression, rRNA metabolic process, tRNA modification, mitochondrial translation.

These processes are enriched in all clusters, showing that their deregulation starts early and is detectable in all the early stages of the disease.

##### RNA molecule and pathway analysis: Insights from MOFA factors

To understand the differences in the pathways and molecules that led to the clusters observed we extracted the top representative pathways and molecules of the MOFA factors. The heatmaps in Fig. 5b show the GSEA normalized enrichment score (NES) of the top representative pathways for each factor for each cluster. Briefly, the score can be interpreted as the correlation of the enriched term with the gene list, and thus higher scores (positive or negative) correlate with genes with higher magnitudes of log2FoldChange (log2FC). SM and IM showed increased NES in pathways related to natural killer cell activation, regulation of leukocyte mediated cytotoxicity, cell killing (pathways enriched in Factor 13) and inflammatory response (Factor 28). The positive NES reflects enrichment at the top of the ranked gene list, ie. the upregulated genes. We observe a decrease in NES in SM in B-cell activation and proliferation, which might be influenced by the reduced peripheral blood B-cell composition noted earlier.

**Fig. 4:**
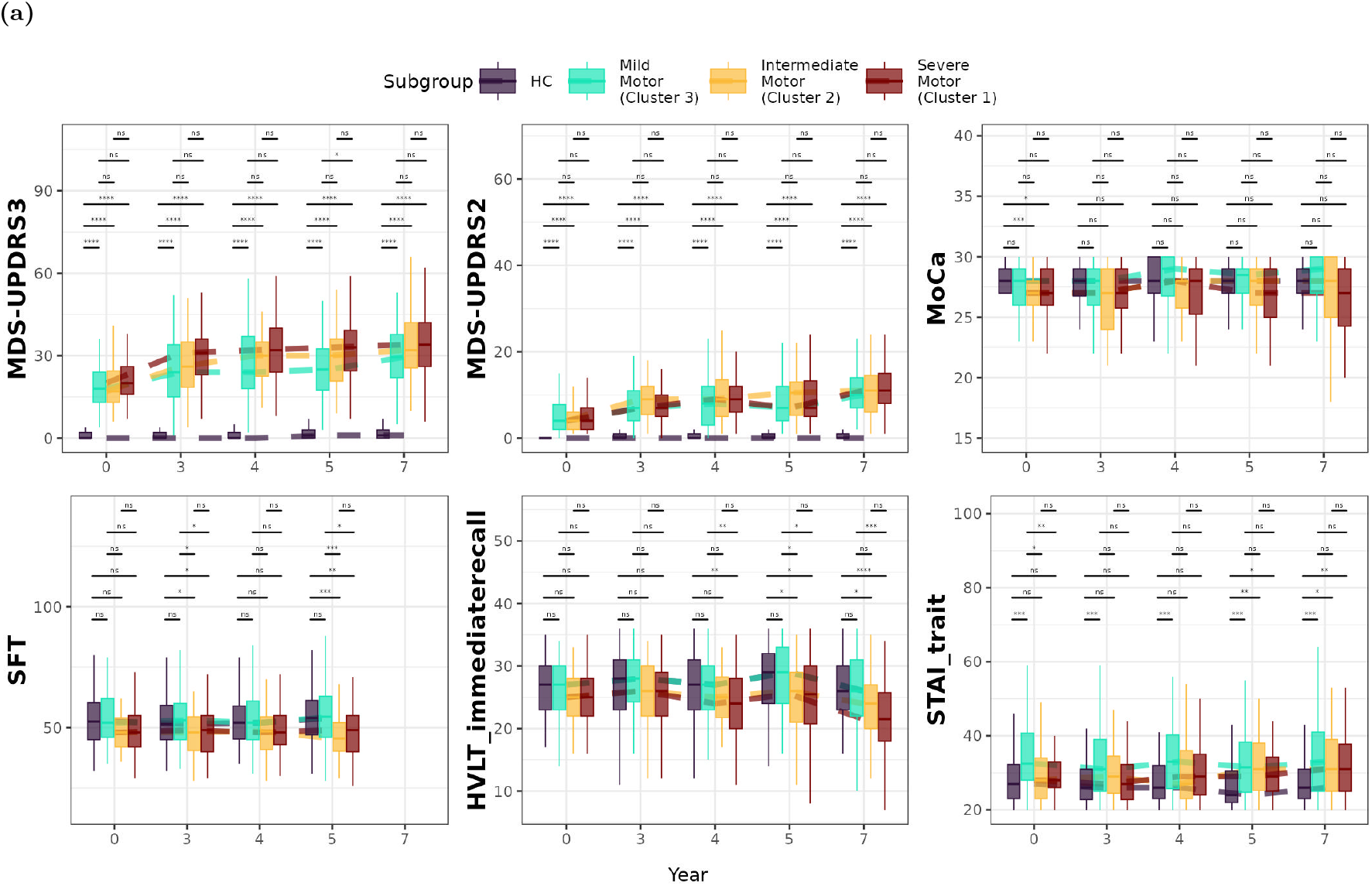
Distribution of clinical scores and markers for clusters obtained using factors correlating with motor severity (MDS-UPDRS3). The boxplot’s middle line indicates the median, and the upper and lower lines indicate the interquartile range. The pairwise differences were tested with the Wilcoxon signed rank test, and the asterisks indicate the statistical p-value (\**p* < 0.05, \*\**p* < 0.01, \*\*\**p* < 0.001, \*\*\*\**p* < 0.0001, n.s. not significant)

**Fig. 5:**
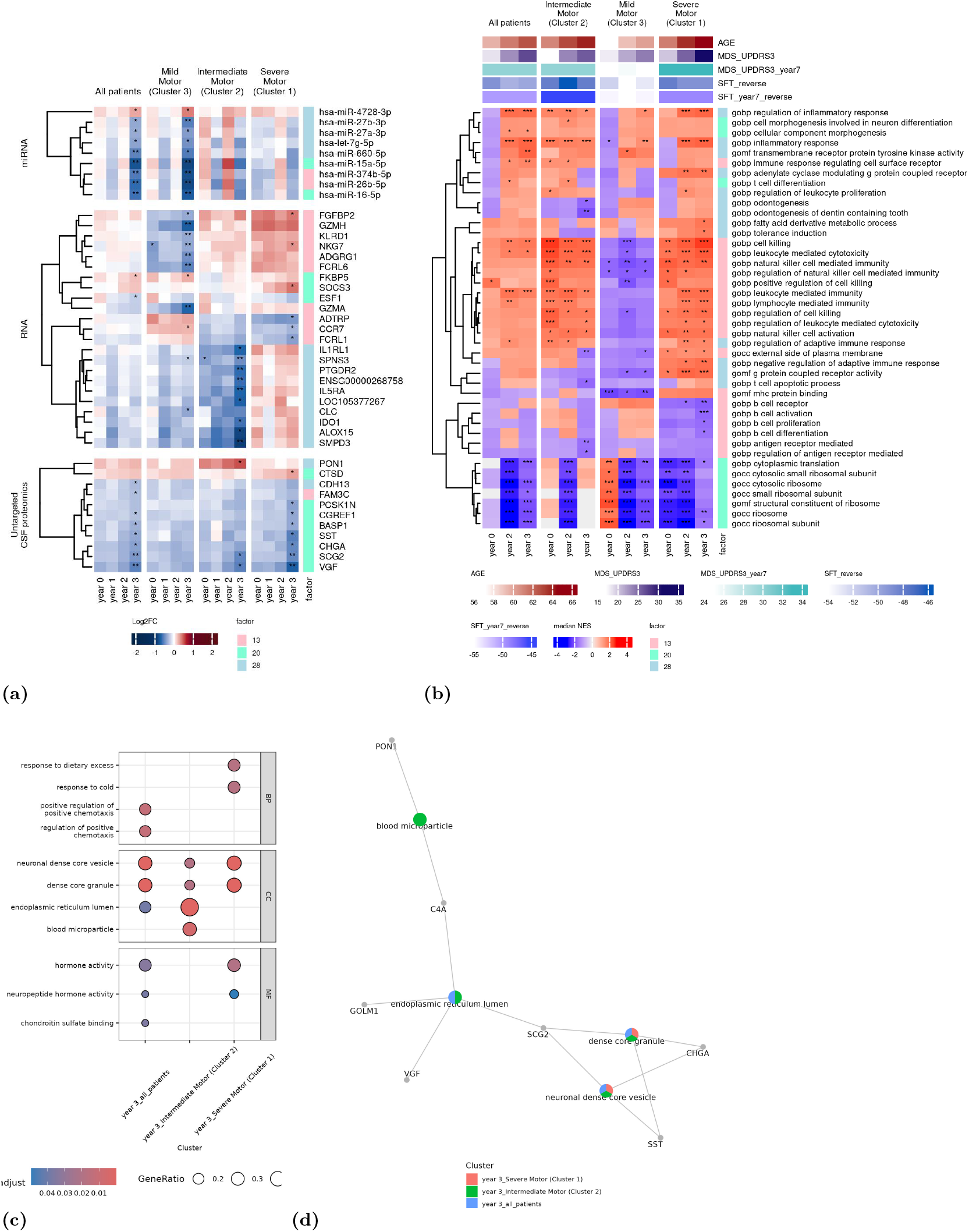
Molecular Signatures of Motor Severity (MDS-UPDRS3) in PD. (a) Differential expression analysis of PD against HC for clusters with differences in MDS-UPDRS3 score, in year 0, 1, 2 and 3. The DE molecules were filtered to show only the highly weighted MOFA molecules. Significantly expressed values compared to HC are marked with *, the p-value is adjusted for age and sex, and Benjamini-Hochberg FDR adjusted for multiple testing. (b) Enrichment analysis of differential expression analysis results against HC using GSEA and the GO database for the RNA modality. Heatmap displays the significantly enriched pathways, filtered for pathways also enriched in the MOFA factors used in the clustering. The coloring represents the normalized enrichment score. (c, d) Overepresentation analysis for the untargeted CSF proteomics. Results are shown only for the clusters that returned significantly enriched pathways compared to HC. The asterisks indicate the statistical p-value (\**p* < 0.05, \*\**p* < 0.01, \*\*\**p* < 0.001, \*\*\*\**p* < 0.0001).

##### Untargeted CSF proteomics Differences relative to Healthy Controls

In CSF proteomics, only the severe and intermediate clusters (SM and IM) showed DE proteins compared to HC. SM showed upregulation of CTSD and downregulation of VGF, CHGA, SCG2, SST, BASP1, CGREF1 and PCSK1N. It is worth noting that these proteins are also in the top 20% highly weighted proteins in the MDS-UPDRS3 mofa factors. Overepresentation analysis of the DE proteins in SM showed among the most highly enriched pathways response to dietary excess (VGF, PCSK1N), protein secretion (SCG2, CHGA, VGF) neuronal dense core vesicle (VGF, SCG2), hormone activity (SST, VGF).

In other omics, untargeted and targeted plasma proteomics and targeted CSF, there were no significantly different molecules which were also highly variable within the top factors, possibly because of the low number of samples with these modalities.

#### 2.2.2 Clustering Patients based on factors correlating with cognitive decline - Semantic Fluency Test (SFT)

##### Overview of clustering approach and clinical correlations

We repeated the clustering of the PD patients using the factors correlated with SFT, which resulted into 3 clusters of n = 65, 48 and 81 patients respectively. Cluster 1 and 2 have a significantly lower SFT score at year 3 and at the last visit recorded (year 5), both compared to Cluster 3 and HC (see Table 2 and Fig. 6). The two clusters also have a worse score on the other cognitive scales of HVLT immediate recall and MoCa. Comparing the other clinical variables we find that Cluster 1 also has higher age and MDS-UPDRS3 than cluster 3. Based on these observations the three clusters were labelled as: Severe Cognitive 1 (SC1), Severe Cognitive 2 (SC2) and Mild Cognitive (MC). SC1 had lower estimations of Bnv and CD4nv cell composition than the MC cluster, and higher CD4mem cells (Supplementary Fig. 3, Supplementary Table 2). SC2 had a larger Bmem cell composition compared to the Mild Cognitive cluster.

**Table 1:**
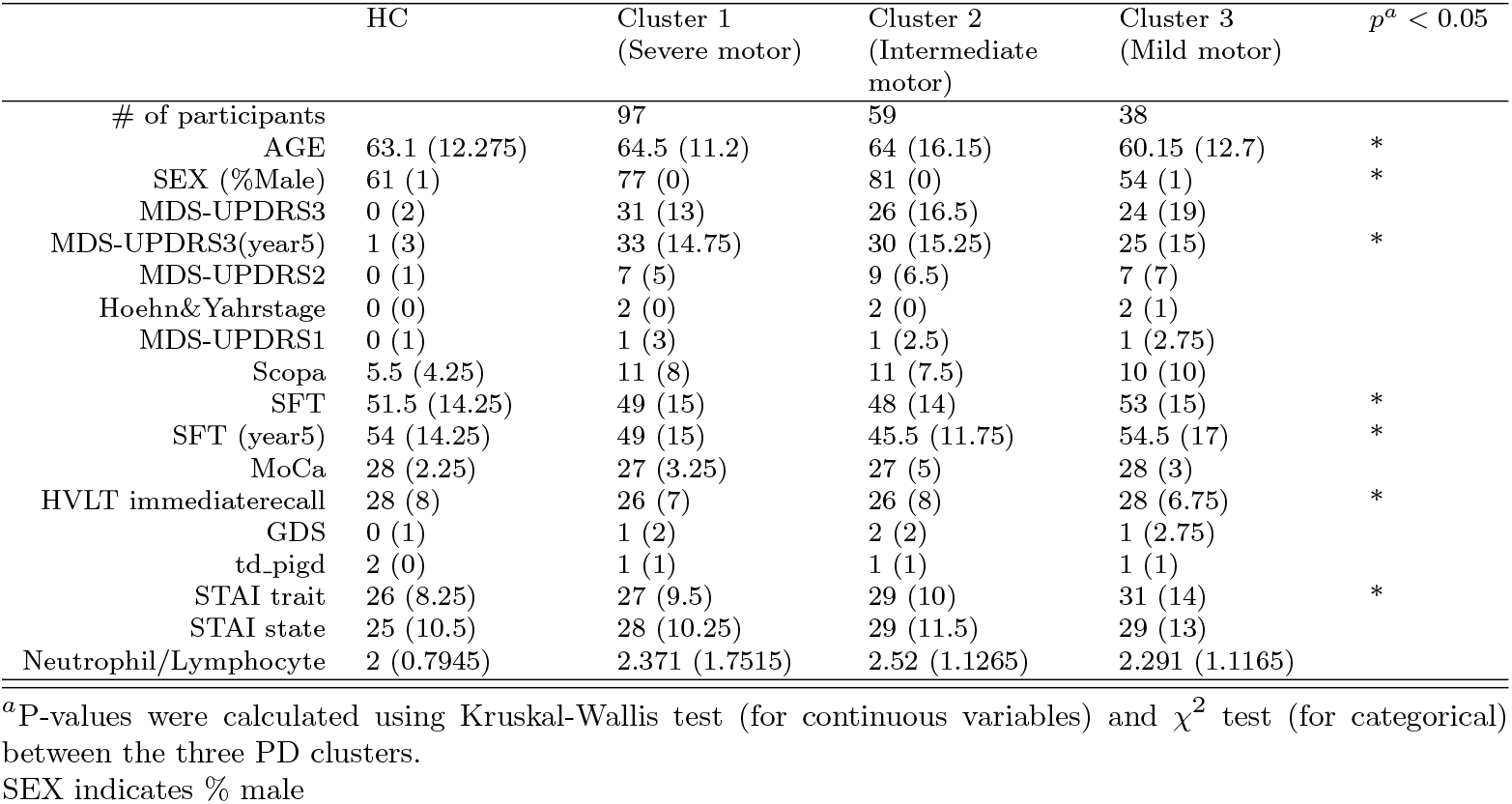
Demographic and clinical variable data summary for the multi-omics clusters with differences in motor severity (MDS-UPDRS3). The table shows the median values and inter-quartile range (IQR) in brackets, for variables in year 3 (unless indicated as a future visit)

**Table 2:**
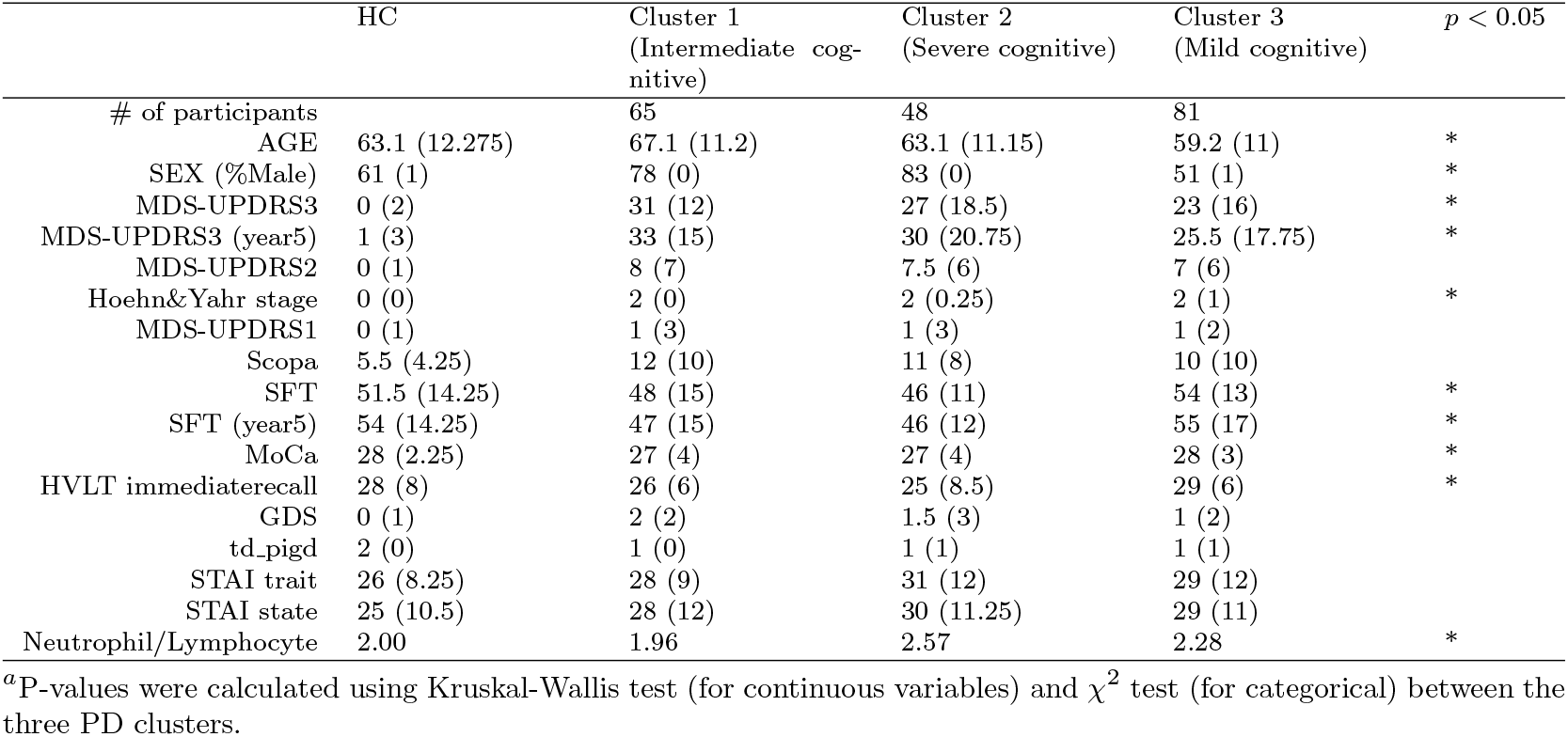
Demographic and clinical variable data summary for the multi-omics clusters with differences in cognitive function (SFT). The table shows the median values and inter-quartile range (IQR) in brackets, for variables in year 3 (unless indicated as a future visit)

**Fig. 6:**
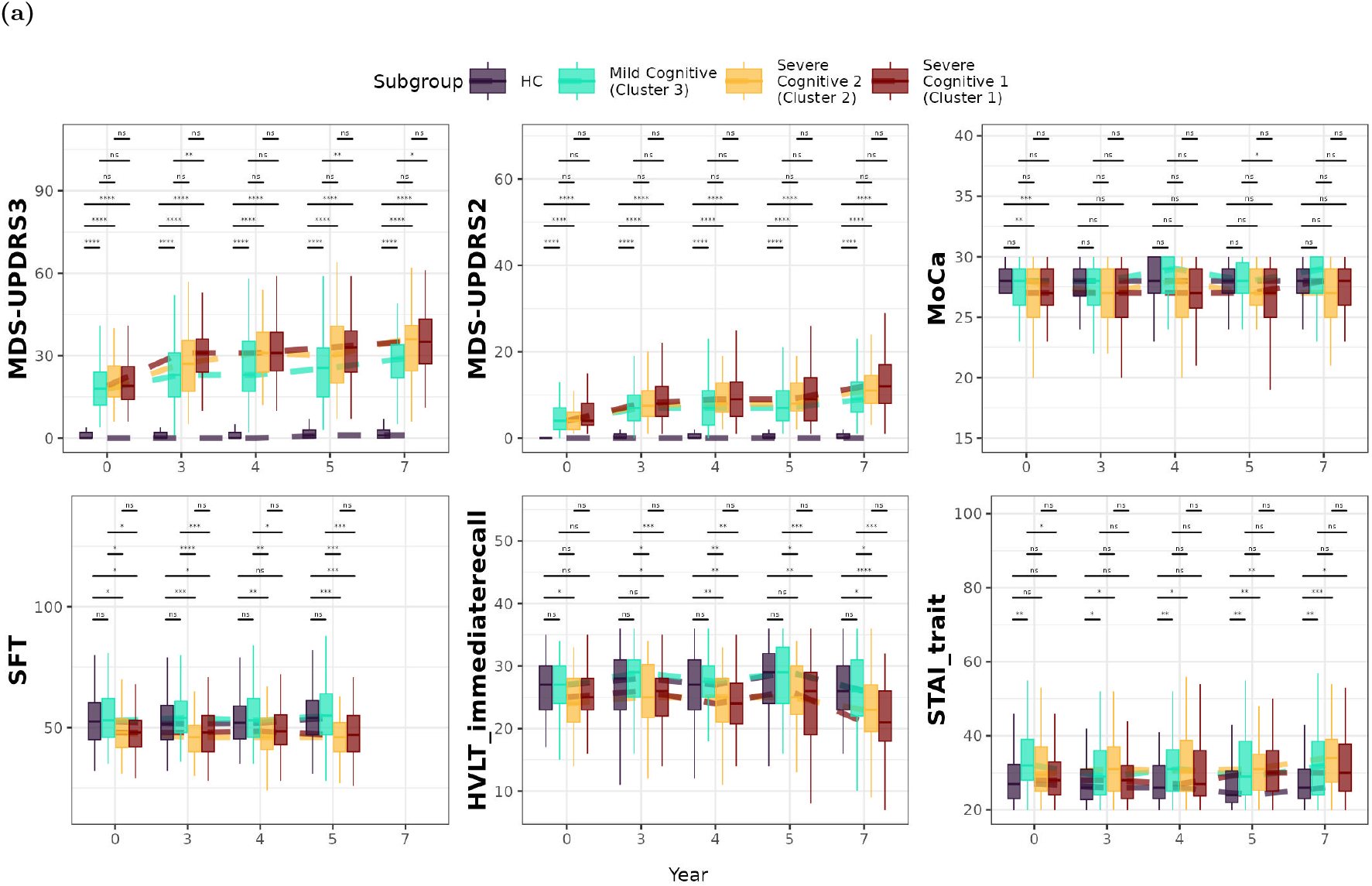
Distributions of clinical scores, covariates and cell types for PD clusters with differences in SFT score. (a) Clinical scores and Covariates. The boxplot’s middle line indicates the median, and the upper and lower lines indicate the interquartile range. The asterisks indicate the statistical p-value (\**p* < 0.05, \*\**p* < 0.01, \*\*\**p* < 0.001, \*\*\*\**p* < 0.0001, n.s. not significant)

From the factors plot (Supplementary Fig. 4) we can see that the severe cognitive 1 cluster is different from the mild cognitive based on factor 13, and the severe cognitive cluster 2 is differentiated from the mild cognitive with factor 16.

##### Top differentially expressed molecular markers and pathways between cognitive decline subgroups

###### RNA Differences relative to Healthy Controls

The most highly enriched pathways include cell killing, natural killer cell-mediated immunity and B-cell receptor signalling. The dysregulation of these pathways in SC1 and SC2 suggests a potential link between immune dysregulation and cognitive severity in PD.

###### RNA molecule and pathway analysis: Insights from MOFA factors

To understand how molecular differences could help explain the differences in the cognitive decline of the PD clusters, we inspected the heatmaps for the top representative pathways and molecules for each factor. Cluster SC1 showed the most differentially expressed pathways against HC which overlapped with the top pathways of the clustering factors (Fig 7b). The top enriched pathways in SC1 included those enriched in factor 13 such as low NES for B-cell activation, and high NES for natural killer cell-mediated immunity. Comparing cluster SC2 to MC, pathway differences from factor 16 included the adenylate cyclase inhibiting G-protein coupled receptor signalling pathway. MC uniquely shows enrichment in pathways of viral genome replication.

**Fig. 7:**
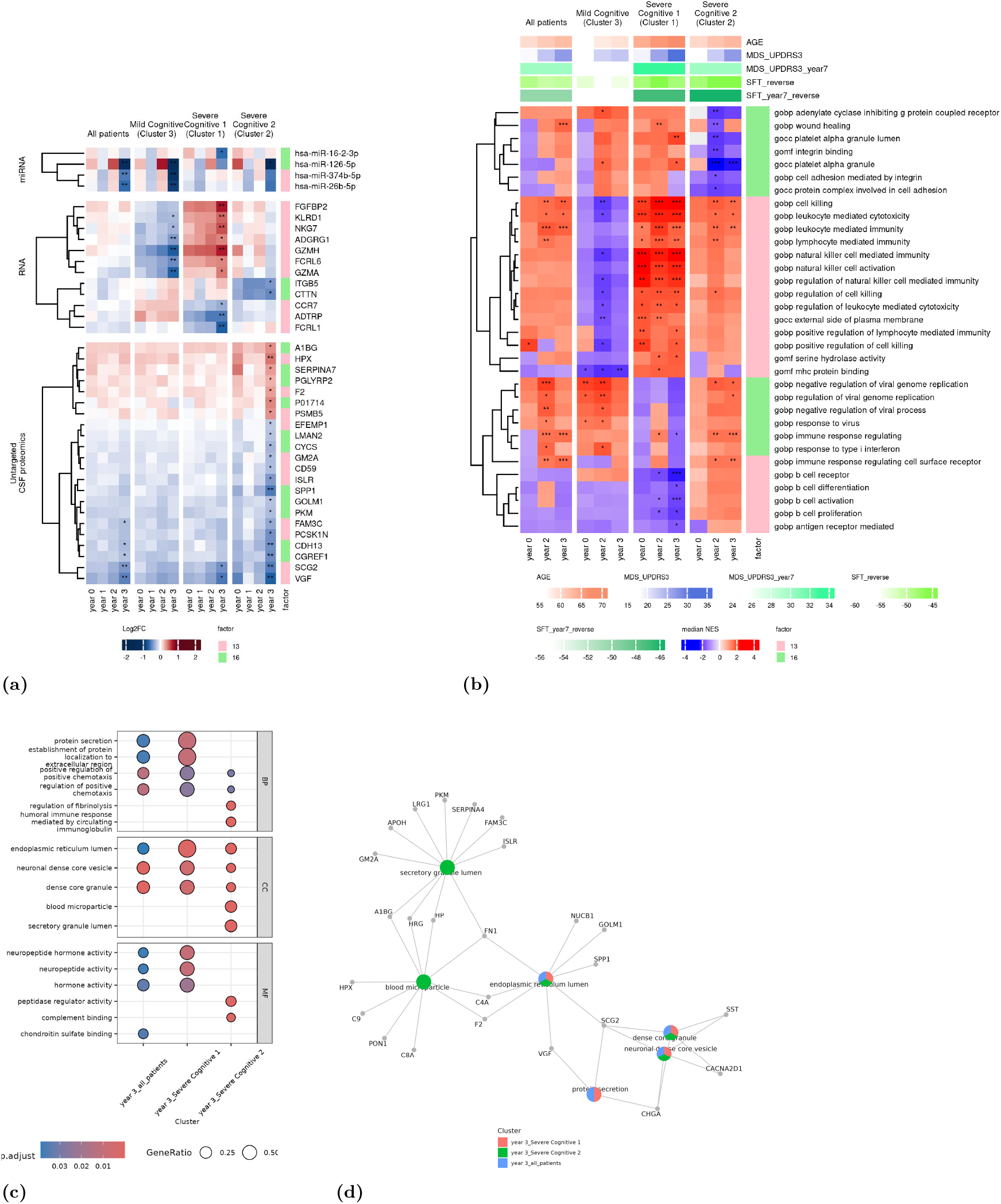
Molecular Signatures of Cognitive Decline (SFT) in PD. (a) Differential expression analysis of PD against HC for clusters with differences in SFT score, in year 0, 1, 2 and 3. The DE molecules were filtered to show only the highly weighted MOFA molecules. The coloring shows the log2FoldChange obtained for each cluster and timepoint. Significantly expressed values compared to HC are marked with *, the p-value is adjusted for age and sex, and Benjamini-Hochberg FDR adjusted for multiple testing. (b) Enrichment analysis of differential expression analysis results against HC using GSEA and the GO database for the RNA modality. Heatmap displays the significantly enriched pathways, filtered for pathways also enriched in the MOFA factors used in the clustering. The coloring represents the normalized enrichment score. (c,d) Overepresentation analysis for the untargeted CSF proteomics. Results are shown only for the clusters that returned significantly enriched pathways compared to HC. The asterisks indicate the statistical p-value (\**p* < 0.05, \*\**p* < 0.01, \*\*\**p* < 0.001, \*\*\*\**p* < 0.0001).

###### Untargeted CSF proteomics differences relative to Healthy Controls

In the proteomics modalities only CSF proteomics revealed differentially expressed proteins for the intermediate and severe cognitive cluster IC, SC2. The logFC values of each cluster at each time point for the top proteins in factors 13 and 16 used in clustering are shown in Fig. 5a. Only cluster SC2 has overlapping differentially expressed proteins with the top proteins in the factors including FAM3C, GOLM1, F2, SCG2, HPX, SERPINA7, PCSK1N, PSMB5, PGLYRP2, LMAN2, PKM allowing us to prioritize these proteins as potential biomarkers for monitoring cognitive dysfunction. Overrepresentation analysis using the differentially expressed proteins showed pathways related to the killing of cells of another organism (F2, C4A, C6), blood microparticle (C8A, C4A, HRG), endopeptidase regulator activity (PCSK1N, SERPINA4, HRG, SERPINA7, C4A), complement binding (C84, CD59C, C4A), secretory granule lumen (SERPINA4, GM2A, LRG1, FAM3C, ISLR, APOH, HP, A1RG, GRG, FN1), endoplasmic reticulum lumen (SPP1, C4A, F2, NUCB1, SPP1, GOLM1) and neuronal dense core vesicle (SST, CACNA2D1, SCG2, CHGA) (see Fig. 7).

###### miRNA differences relative to Healthy Controls

For miRNAs hsa-let-7d-3p, and miR-3157-3p are found to be overexpressed in SC1 while hsa-miR-126b-5p and miR-16-2-3p was underexpressed in SC2. Interestingly, for the miRNA modality, the most differentially expressed (DE) molecules that overlap with the top molecules in factors are in the mild cognitive cluster (MC), and include hsa-miR-26b-5p, hsa-miR-374a-5p and hsa-miR-126-5p.

### 2.3 Computational Drug Repurposing using the CSF proteins

We performed computational drug repurposing targeting the differentially expressed (DE) proteins in CSF, aiming to identify drugs capable of reversing their gene expression. Only proteins with consistent direction in gene expression confirmed in the literature or external datasets were included. We confirmed the direction of gene expression in brain tissue or CSF for three proteins in the SM cluster: BASP1 in [21] and VGF and CGREF1 in the GSE205450 dataset. In the SC2 cluster, seven proteins were confirmed: SCG2, FAM3C, VGF, SPP1, NUCB1, CGREF1, BASP1. VGF, SCG2, CGREF1, FAM3C were confirmed in GSE205450 and the rest in literature: NUCB1 [22], SPP1 [23], BASP1 [21]. The SM cluster proteins confirmed, were were all part of the SC2 cluster and thus drug repurposing was performed once for the seven SC2 proteins. We generated seven lists of drugs (one per gene) and filtered them for compounds that can reverse the expression of at least 50% of these genes, which resulted in a list of 10 drugs: BRD-K19362120, BRD-K46267071, BRD-K86574132, KM-00927, SEW-05685, WH-4025, enzastaurin, fenretinide, ispinesib and verrucarin-a. To understand their potential therapeutic relevance, we retrieved the mechanism of action in the Drug Repurposing Hub [24], identifying known functions for three drugs: Enzaustarin, a PKC inhibitor, ispinesib, a kinesin inhibitor and fenretinide, an apoptosis stimulant and retinoid receptor agonist.

## 3 Discussion

This study aimed to detect molecular factors associated with the progression of idiopathic Parkinson’s disease. Unlike previous studies that have used a selective approach, our study has taken a more exploratory, data-driven approach by using multi-omics factor analysis to identify patterns of biomarkers associated with disease state and severity. The results of the study revealed three molecular factors related to motor severity (measured by MDS-UPDRS3), and two factors associated with cognitive decline (measured by the Semantic Fluency Test, SFT). The distinct factors related to motor and cognitive scores, indicate that the progression of these symptoms in PD follow separate trajectories. We thus clustered the patients with two different sets of factors to separate the patients into groups of different severities of the scales of motor and cognitive dysfunction.

### Pathways and molecules associated with motor dysfunction differences

Using k-means clustering of the molecular factors we classified patients into three motor severity clusters: Severe Motor (SM), Intermediate Motor (IM) and Mild Motor (MM). These subgroups exhibited distinct molecular features, which could be useful for detecting these stages in clinical settings. The produced subgroups showed differences in age and sex, with the two most severe subgroups having higher age and a higher male predominance which aligns with known risk progression factors in PD at least in the first years after diagnosis [25]. The higher median MDS-UPDRS3 in SM and IM at year 3 and also in the latest year recorded (year 7) underscores the progressive motor impairment in these subgroups.

Several molecular processes detected in this study had been previously associated with PD pathogenesis, and more specifically associated with the measured clinical scores or the progression of PD. In RNA sequencing data, the severe motor subgroup of patients showed differences in natural killer cells and T-cell immunity, the expression of B-cell-related genes, and depletion in CD4 naive cells compared to the mild motor subgroup, likely reflecting peripheral immune dysregulation observed in severe motor phenotypes. Previous studies showed that enhanced immune response of B lymphocytes, abnormal distribution of B cell subsets [26] and specifically the serum levels of B cell-associated proteins correlated negatively with MDS-UPDRS3. Single cell RNA-sequencing data from fresh blood of early-to-mid PD patients, reported similar results [27], and flow cytometry confirmed depletion of CD4 naive cells in peripheral blood [28]. The increase in NK cell activity was previously reported in PD and correlated with severity. Their role is not established yet but in-vitro experiments indicated a potentially protective effect of these cells in PD [29]. Differential expression analysis of RNA sequencing identified a large number of molecules, which, after adjusting for immune cell composition as covariates were greatly reduced. This indicates that immune cell heterogeneity is a primary driver of whole-blood RNA variability.

The CSF proteomics analysis revealed the neurosecretory protein VGF and Chromogranin A (CHGA) which belong to the granin family are down-regulated in the severe motor clusters compared to healthy controls. These proteins have numerous neuronal functions, including regulation of neuroendocrine pathways, microglial activation, and synaptic plasticity, and were previously found to decrease in PD [30]. Since granins are co-stored in dense core vesicles and can regulate levels of catecholamines such as dopamine, the decrease of their abundance in clinically severe PD groups could co-occur with the progression of catecholaminergic deficit in PD. Other deregulated proteins in the severe motor clusters was SCG2, which is connected to the endoplasmic reticulum (ER), showing ER dysfunction as the disease progresses to later stages. The ER is responsible for protein folding and dysfunction can lead to ER stress. Misfolded *α*-synuclein accumulates and overwhelms the ER leading leads to chronic ER stress [31]. BASP1 regulates actin dynamics and presynaptic vesicle cycling at axon terminals, thereby facilitating axonal growth, regeneration, and plasticity [21]. Dysregulation of BASP1 disrupts axonal integrity potentially impairing motor control in PD.

### Pathways and molecules associated with differences in cognitive decline

The cognitive severity was assessed using the Semantic Fluency Test score. Semantic fluency relies on the efficient retrieval of words from memory, a process that involves various brain regions, including the frontal and temporal lobes. The severe cognitive clusters showed similar biological pathways to severe motor clusters in blood RNA, including pathways associated with NK cells, T-cell immunity and B-cell gene expression differences compared to the mild cognitive subgroup.

In the severe cognitive clusters, proteins F2, C6, C4A were de-regulated, representing pathways like the blood microparticle and complement binding. This could indicate a stage where complement proteins are either produced by the brain or infiltrate from the periphery [32]. Fibrinogen and complement Factor H were previously associated with neuropsychological test scores and cognitive impairement in PD [33]. Other proteins like SERPINA7, GM2A, and FAM3C are involved in secretory granule function, potentially implicating deficits in neuronal vesicle trafficking, which is critical for neurotransmitter release. GM2A is similarly elevated in AD [34]. Previous studies on Huntington’s disease, showed that selecting patients with evidence of complement system activation, is more effective, since they respond significantly better to anti-complement drugs, compared to patients without evidence of complement [35]. This shows that the detection of subgroups with differences in complement system in PD could also guide the selection of patients that should be recruited to these trials. The enrichment of neuronal dense core vesicle (DCV) pathway proteins (CHGA, SST, CACNA2D1 and SCG2) which are important both for vesicle formation and their release at the synapse indicates a shared pathophysiology with motor dysfunction clusters. These results highlight the relevance of vesicular trafficking in both motor and cognitive impairments in PD which has a key role in the etiology of PD [36]. DCV carry critical neurotransmitters like dopamine, serotonin and neuropeptides which modulate synaptic plasticity. Dysfunction of DCV and disruption of vesicular homeostasis can interfere with the balance of neurotransmitters that contribute to motor and non-motor symptoms in PD.

### Potential of identified drugs for Parkinson’s Disease

The identified drugs were selected based on their ability to reverse the gene expression of key differentially expressed proteins in the CSF and brain tissue. Among the ten identified drugs, fenretinide and enzaustarin have been previously studied in neurological disease, although not specifically in PD. Fenretinide has been shown to target several pathways in neurological disease including oxidative stress and the inflammatory response [37], making it a promising molecule for PD. Enzastaurin, a protein kinase (PKC) inhibitor, was shown to ameliorate neuroinflammation, axonal damage and clinical symptoms of paralysis in experimental autoimmune encephalomyelitis mice [38]. Ispinesib may impact the lysosomal pathways and intracellular trafficking [39], which could be relevant for protein aggregation and transport in PD. KM-00927 is predicted to be an HDAC inhibitor [40]. HDACs are involved in several mechanisms related to PD, including regulation of neurogenesis, proteolysis and immune responses through mechanisms like epithelium and endothelial migration. HDAC has also been identified within Lewy bodies. Notably, inhibition of HDAC6 has shown protection of dopaminergic neurons from *α*-synuclein toxicity in [41]. Data on the other compounds are limited but their potential mechanisms might be beneficial to PD. BRD-K86574132 is predicted to be a PI3K inhibitor, and was previously recommended for Hepatocellular Carcinoma based on its potential to target oxidative stress-related genes [42]. Oxidative stress is also central to PD pathology and targeting these genes with BRD-K86574132 could improve disease symptoms. Verrucarin-a, an inhibitor of protein synthesis [43] was found to have apoptosis-inducing activity, but was previously found to be toxic for use in animal models. Nevertheless, insights into the mechanism of action might direct the search for similar drugs with safer application.

Overall, the drugs identified, target key pathways associated with Parkinson’s disease pathology, including neuroinflammation, oxidative stress, lysosomal pathways and intracellular trafficking making them potential candidates for the disease. The identification of drugs with no prior PD studies underscores the utility of multi-omics guided drug repurposing.

### Limitations and Further Work

Our study focused on early-stage Parkinson’s disease (PD) patients, using molecular data from individuals within three years of diagnosis. At this stage, clinical changes may still be subtle, leading to weaker correlations between molecular mechanisms and disease severity. We anticipate that repetition of the analysis with more progressed patients will identify greater and higher correlations of the molecular mechanisms with the clinical image. One key limitation is the use of the maximum value of the MDS-UPDRS3 score, to approximate motor severity, which is affected by symptomatic drug medication, leading to fluctuations throughout the day. This is a general complication of assessing the severity of Parkinson’s since the symptomatic medication can cover the molecular severity of the disease. In future works we would like to take into account the effect of medication, to obtain an accurate representation of the clinical severity. Finally, variations in blood cell type compositions create confounding effects for the molecular states we observe, and makes it difficult to draw mechanistic conclusions across omics layers. Single-cell technologies could provide more detailed insights into the mechanisms related to severity by allowing cell type-specific measurements and cellular multi-omic layer connections. A similar multi-omics analysis approach as the one presented in this work, applied at single cell level is among our future plans where there will be proper available data to work with.

## 4 Conclusion

Our findings provide a multidimensional view of the severity of PD through the integration of RNA expression and proteomics datasets from multiple tissues and clinical scores. The identification of immune and vesicular pathways across motor and cognitive clusters suggests that PD severity is driven by complex, intersecting biological processes. Overall, several markers prioritized by the multi-factor analysis have established associations with the pathology of Parkinson’s disease and are strong candidates to serve as monitoring markers for the progression of motor and cognitive dysfunction. Future work should aim to validate these findings in larger cohorts. Finally, investigating therapeutic strategies targeting immune modulation and dense core vesicle pathways could help to mitigate disease progression in the early stages of PD.

## Supporting information

Supplementary Dataset 1 - Differential Expression Analysis of PD clusters associated with MDS-UPDRS3

Supplementary Dataset 2 - Differential Expression Analysis of PD clusters associated with SFT

## Acknowledgements

This study used data from the PPMI (obtained from PPMI database at www.ppmi-info.org/access-data-specimens/download-data). PPMI—a public-private partnership—is funded by the Michael J. Fox Foundation for Parkinson’s Research and funding partners, including 4D Pharma, Abbvie, AcureX, Allergan, Amathus Therapeutics, Aligning Science Across Parkinson’s, AskBio, Avid Radiopharmaceuticals, BIAL, BioArctic, Biogen, Biohaven, BioLegend, BlueRock Therapeutics, Bristol-Myers Squibb, Calico Labs, Capsida Biotherapeutics, Celgene, Cerevel Therapeutics, Coave Therapeutics, DaCapo Brainscience, Denali, Edmond J. Safra Foundation, Eli Lilly, Gain Therapeutics, GE HealthCare, Genentech, GlaxoSmithKline plc (GSK), Golub Capital, Handl Therapeutics, Insitro, Janssen Neuroscience, Jazz Pharmaceuticals, Lundbeck, Merck, Meso Scale Discovery, Mission Therapeutics, Neurocrine Biosciences, Neuropore, Pfizer, Piramal, Prevail Therapeutics, Roche, Sanofi, Servier, Sun Pharma Advanced Research Company, Takeda, Teva, UCB, Vanqua Bio, Verily, Voyager Therapeutics, the Weston Family Foundation and Yumanity Therapeutics. For up-to-date information on the study, visit www.ppmi-info.org. This work was supported by TELETHON Cyprus.

Supplementary Information is available for this paper.

## Methods

### Multi-omics data

We utilize the Parkinson’s Progression Marker Initiative (PPMI) [19] dataset, a longitudinal Parkinson’s Disease (PD) cohort, which includes clinical and multi-omics data for PD patients. Data used in the preparation of this article were obtained on 11 March 2024 from the PPMI database (www.ppmi-info.org/access-data-specimens/download-data), RRID:SCR 006431. For up-to-date information on the study, visit www.ppmi-info.org. We used omics (blood RNA, blood miRNA, and plasma and CSF proteomics) from patients (idiopathic PD) and healthy controls across four-time points, years 0, 1, 2 and 3 which correspond to the PPMI Visit codes: BL, V04, V06 and V08. We selected the RNA and miRNA data from the PPMI RNA sequencing project, from which we used the 423 de novo idiopathic PD patients and 196 healthy controls (HC). The RNA isolation was done at the biorepository at Indiana University and then sent to HudsonAlpha Biotechnnology for library preparation and sequencing. The total RNA was sequenced in a strand-specific manner to a depth of 100 million read pairs per sample. More details into the methods of data processing and analysis of this project are available at https://ida.loni.usc.edu/web/ppmi-rnaseq-app. We also downloaded the data produced on targeted CSF and plasma proteomics from the project 9000, and the untargeted proteomics from project 177, details can be found at https://www.ppmi-info.org/sites/default/files/docs/PPMI%20Data%20User%20Guide.pdf. Project 9000 is a longitudinal Olink proteomics study of de-novo PD and non-genetic prodromal participants age 60 or older with REM sleep behavior disorder (RBD) or hyposmia and Dopamine Transporter (DAT) imaging deficit, with repeat sampling over 4 years. The data were provided in Olink NPX units, a normalized arbitrary unit derived from sequence counts. Project 177 is a longitudinal LC-MS proteomics study from 482 participants from CSF and 179 from blood plasma. The LC-MS data was already processed through the OpenSwath pipeline, run through mapDIA and batch corrected.

### Clinical scores and biochemical markers

At the time of this project, the clinical data were collected until year 10 after diagnosis. The clinical measurements used, include motor and non-motor (cognitive, neurobehavioral, neuropsychological, autonomic, olfaction, sleep) data. Namely, the scores used were MDS-Unified Parkinson’s Disease Rating Scale (MDS-UPDRS), NeuroBehaviour, Cognitive testing (eg. MoCa), Autonomic Testing (SCOPA), the Semantic Fluency test (SFT), the State-Trait Anxiety inventory (STAI), the Geriatric Depression Scale (GDS) and Hopkins Verbal Learning Test (HVLT) (see Table 3 for descriptions). At baseline, all participants with PD were free of dopamine-related medications. Use of medications for PD was recorded at the 6- and 12-month visits, and is expressed as levodopa equivalent dose (LEDD). In addition, several biochemical measurements including CSF amyloid-beta1–42 (*Aβ*_1−42_), alpha-synuclein (a-syn), total tau (t-tau) and tau phosphorylated at Thr181 (p-tau), and the ratio of *Aβ*_1−42_*/α*-syn (ab asyn), and tau/*α*-synuclein (tau asyn) are available for all patients at each time point.

**Table 3:**
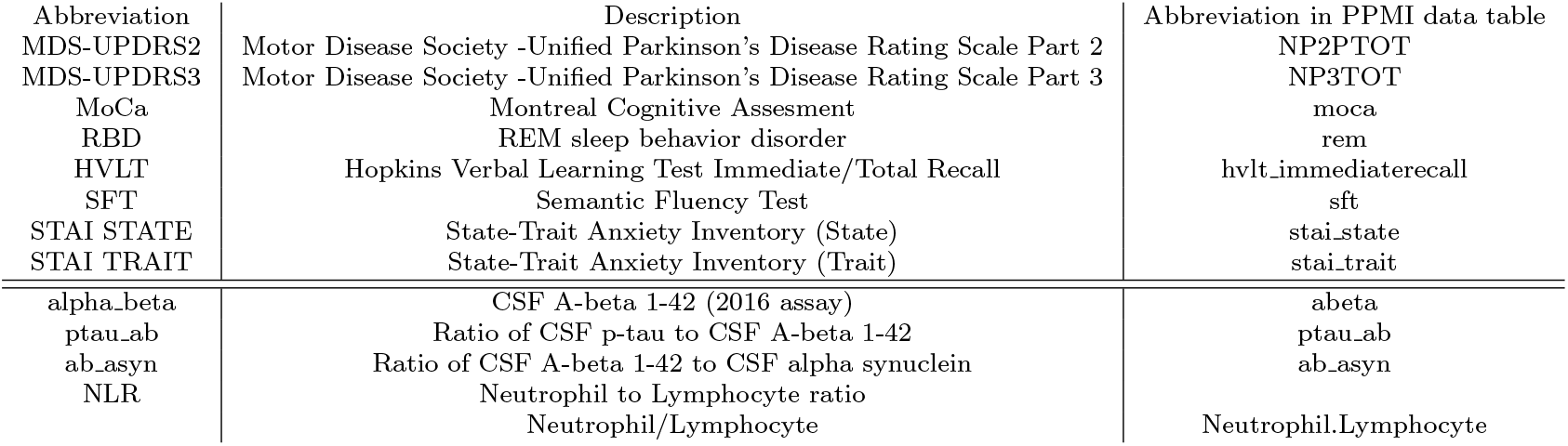
Clinical scores abbreviations and full description and abbreviations used in the PPMI database.

### Estimation and correction for cell type composition in blood

The cell type composition of Whole Blood RNA sequencing samples, was estimated using the package `granulator v1.10.0` [44] which uses a set of published algorithms and reference datasets for the estimations. In the analysis, the results of the algorithm `qprog` were used because it showed the highest correlation with the measured Neutrophil percentage in PPMI patients.

### Batch correction of the input data

The RNA sequencing (small and long RNA) data were adjusted using a Bayes Linear Model from the `WGCNA v1.72.5` [45] package before factor analysis. The data first underwent variance transformation stabilization. Then the correction was performed using `usable bases`, `plate` as the covariates to remove, and the covariates to preserve were: `disease status, age` and `sex`.

### Multi-Omics factor analysis

For factor analysis, we used molecular data from year 3 after diagnosis (PPMI Visit V08) to identify factors associated with disease severity. By this time point, the molecular profile of Parkinson’s disease is more fully developed, and patients have established distinct severity states. This selection ensures that the factors detected are representative of stable molecular and clinical differences, minimizing confounding variability associated with early disease stages or fluctuating presentation of symptoms. Factor analysis was performed using the package `MOFA2` with 34 factors using six data modalities the (a) RNA and (b) miRNA sequencing data (after variance transformation and batch correction), the targeted proteomics dataset from (c) plasma and (d) CSF, and untargeted proteomics from (e) plasma and (f) CSF. To decide on the number of factors we tried various values within the interval of [20-50] and chose the value where the pearson correlation of factors with the *cohort* value (which indicates 1 for PD and 2 for HC) would be maximized. Dimensionality reduction methods work well with an input of the most variable features, usually recommended to be the top 25% for RNA sequencing data analysis (eg. [46]). The most variable features were obtained by ordering them by their standard deviation across samples. For multi-omics analysis, we tried to balance the number of features for each omics dataset, and thus because the RNA modality is significantly larger than proteomics and miRNAs, we used the following numbers for each: the 20% most variable RNA features (resulting in 2161 transcripts), 50% miRNAs (314 transcripts) and 90% proteomics (167, 262, 730, 732 proteins for untargeted plasma, untargeted CSF, targeted plasma and targeted CSF proteomics). The correlations of factors with clinical scores and other covariates were calculated using Pearson correlation using `corr.test` from the `psych` [47] package.

### Molecular subtypes and clinical associations

The clustering of patient samples was performed using the k-means algorithm and MOFA factor values as input. We used factors that were significant predictors (p<0.05) of the (a) MDS-UPDRS3 and (b) SFT score using linear regression analysis after adjusting for confounding variables age, sex and LEDD. To assess the differences of the clinical variables between the clusters, we performed the *χ*^2^ test for discrete variables and the Kruskal-Wallis test for continuous variables. For the pairwise associations we used the Wilcoxon signed rank test.

### DE analysis

For the DE analysis for small/ long RNA-seq, linear modelling and empirical Bayes moderation was done with DESeq2 [48], using Sex, Age, Plate and Usable bases as covariates in the design formula in addition to disease status, and where stated, the blood cell type composition, since they were shown to be the primary sources of gene and transcript variation [49]. Pre-processing was done to filter lowly expressed transcripts using `filterByExpression` and `EstimateSizeFactors` from the DESeq2 package. The DE analysis of the proteomics datasets was performed using Limma [50], and were corrected for Age and Sex. For all modalities, we considered statistically differentially expressed (DE) any features with a p-adjusted < 0.05 and |*log*2*FC*| ≥ 0.1.

### Enrichment analysis

Gene ontology term and pathway enrichment analyses were performed using several common functional annotation databases (GO-biological processes (BP), cellular component (CC) and molecular functions (MF)) using the tool `ClusterProfiler v4.10.0` [51] to obtain enriched pathways in PD patient clusters compared to HC. For the RNA modality, enrichment analysis was done with Gene-set Enrichment analysis (GSEA) using the output from DESeq2, and the genes were ranked by *log*2*FC*. The enrichment results for the RNA modality using GSEA are presented with the Normalized Enrichment Score (NES) of each term [52]. For proteomics, over-representation analysis (ORA) was performed using the significant proteins obtained with limma with p-adjusted< 0.05. The enrichment result plots were obtained using the function `compareCluster` from the `ClusterProfiler` package. For MOFA factors, enrichment analysis was done using Principal Component Gene Set Enrichment `PCGSE` [53] using the function `runEnrichmentAnalysis` in the MOFA package.

### Computational Drug Repurposing

To confirm the consistency of the direction of gene expression of DE CSF proteins, we reviewed the literature and analyzed expression data from the GSE205450 dataset [54] (transcriptomics data of human post-mortem PD striatum). These validated genes were input into the Gene Budger tool [55] to obtain the drugs that reverse the transcriptomic profile, following the pipeline in [56]. Briefly, we received a list of drugs that significantly modulate the expression of each of the genes with a |*log*2*FC*| ≥ 1. We excluded the drugs that could both reverse and enhance the specific gene to discard those with non-specific effects. We then filtered the lists by the drugs that could alter at least 50% of the input genes.

