## Supplementary File for "Multi-omics dissection of Parkinson’s patients in subgroups associated with motor and cognitive severity"

### Supplementary Information

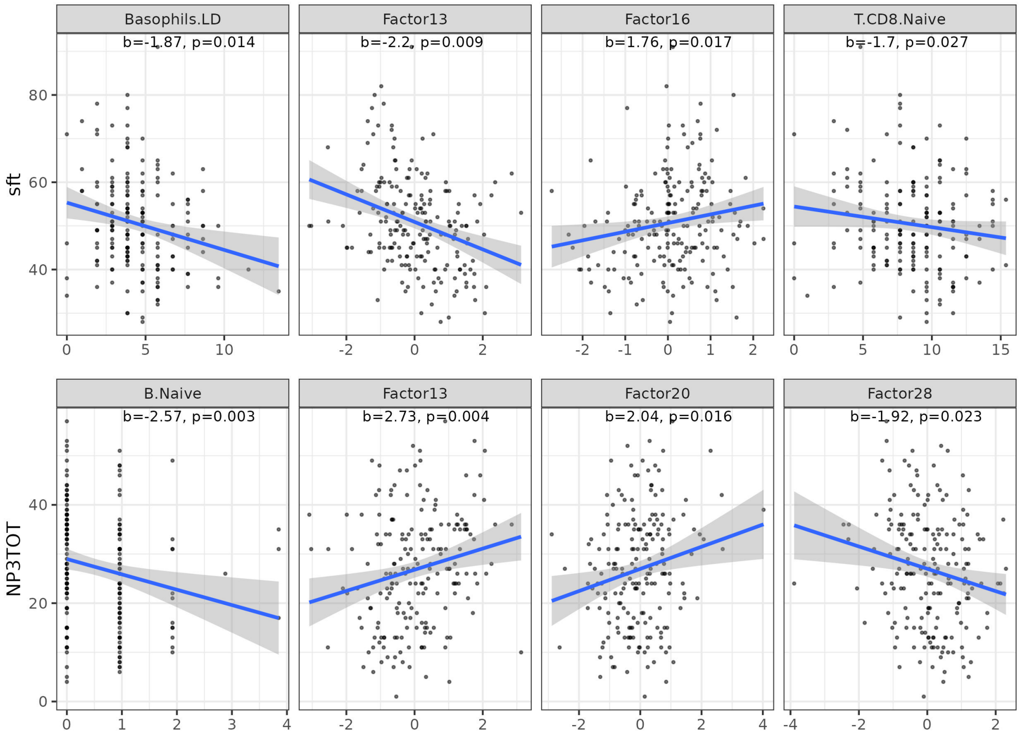


**Supplementary Figure 1: Linear fit for Factors and cell type covariates against clinical scores**

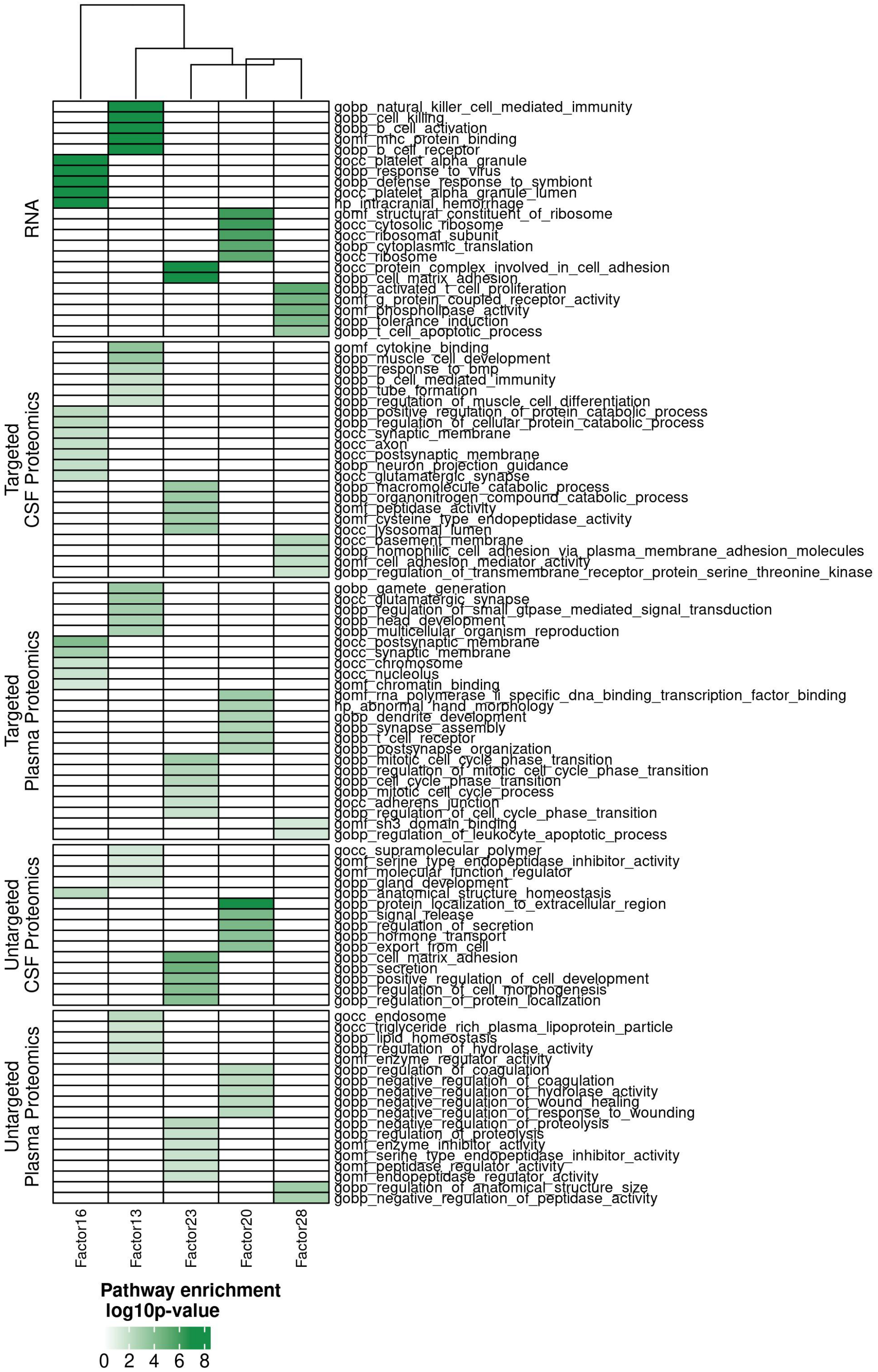


**Supplementary Figure 2**: **Factor annotations: Pathways** Top pathways of each factor using Principal Component Gene-set enrichment analysis (PCGSE). The coloring represents the significance of pathway enrichment in log10(p-value).

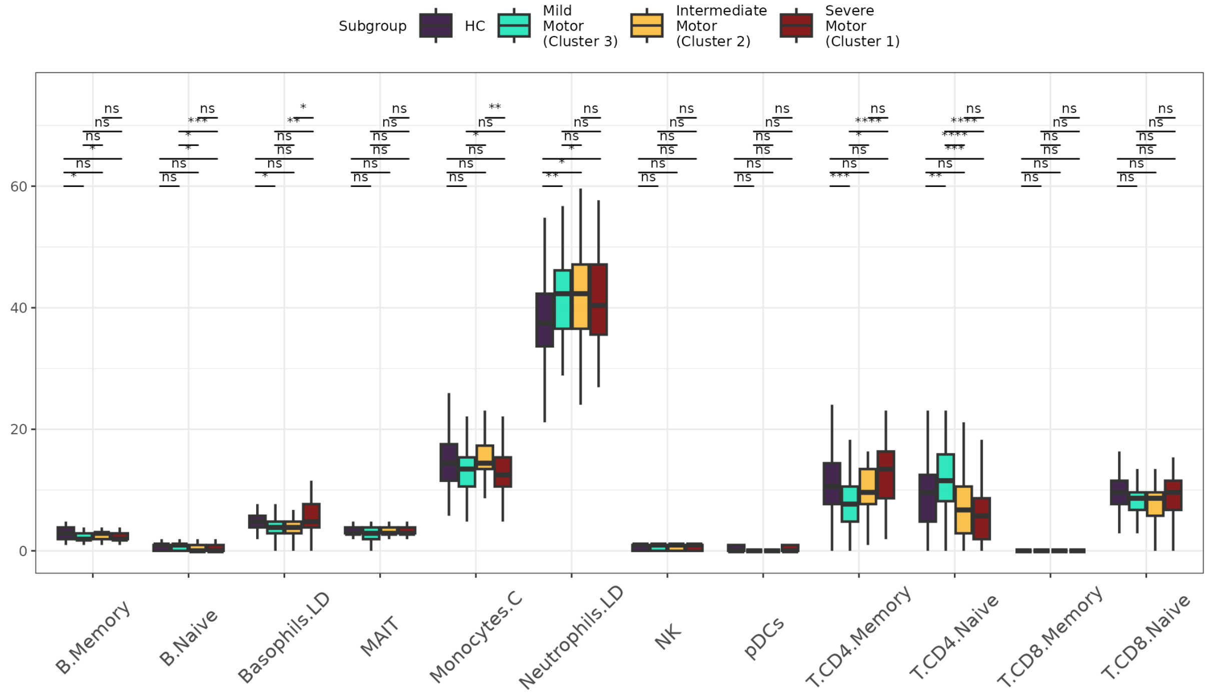


**(a)**

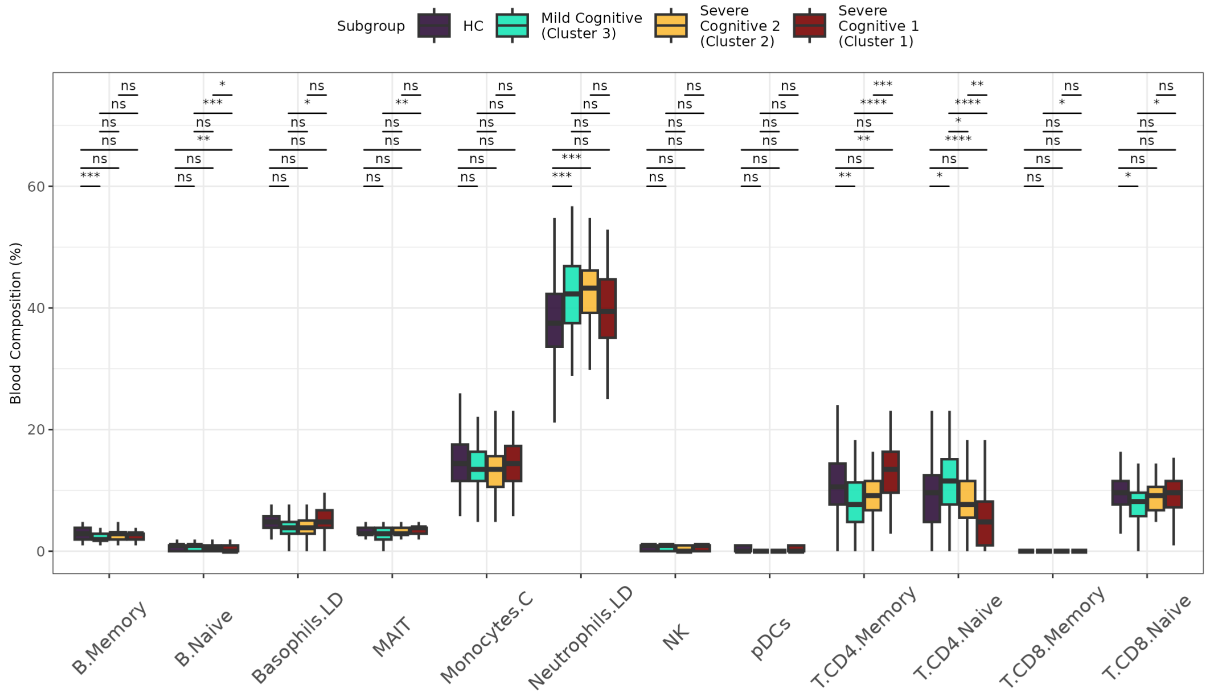


**(b)**

**Supplementary Figure 3**: Median proportion (%) of cell types for PD patient clusters with differences in (a) MDS-UPDRS3 and (b) SFT. The boxplot's middle line indicates the median, and the upper and lower lines indicate the interquartile range. The pairwise differences were tested with the Wilcoxon signed rank test, and the asterisks indicate the statistical p-value (*$p < 0.05$, **$p < 0.01$, ***$p < 0.001$, ****$p < 0.0001$, n.s. not significant)

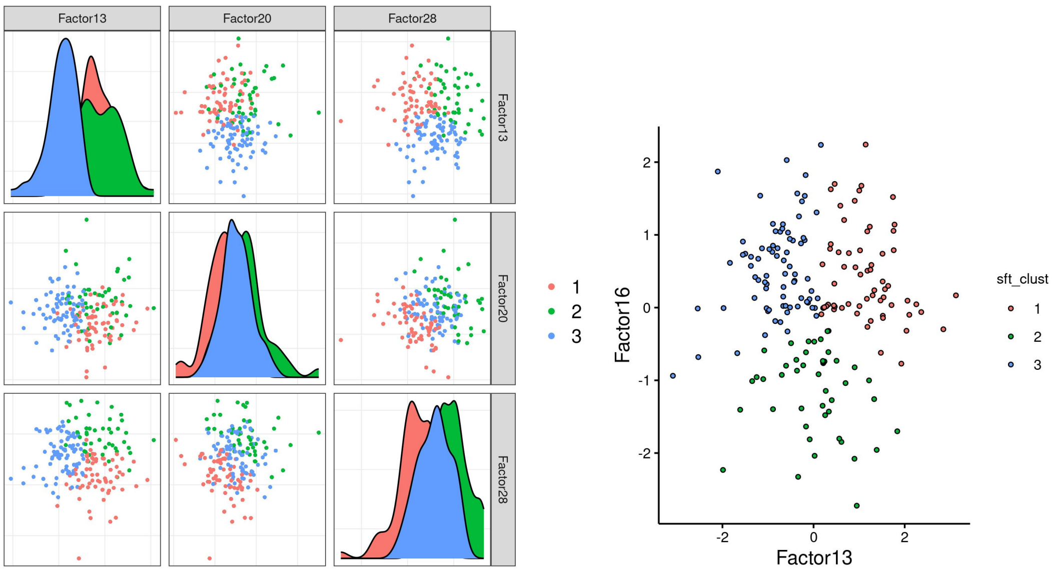


**Supplementary Figure 4**: Mofa factor distribution for each PD cluster for the factors used in clustering

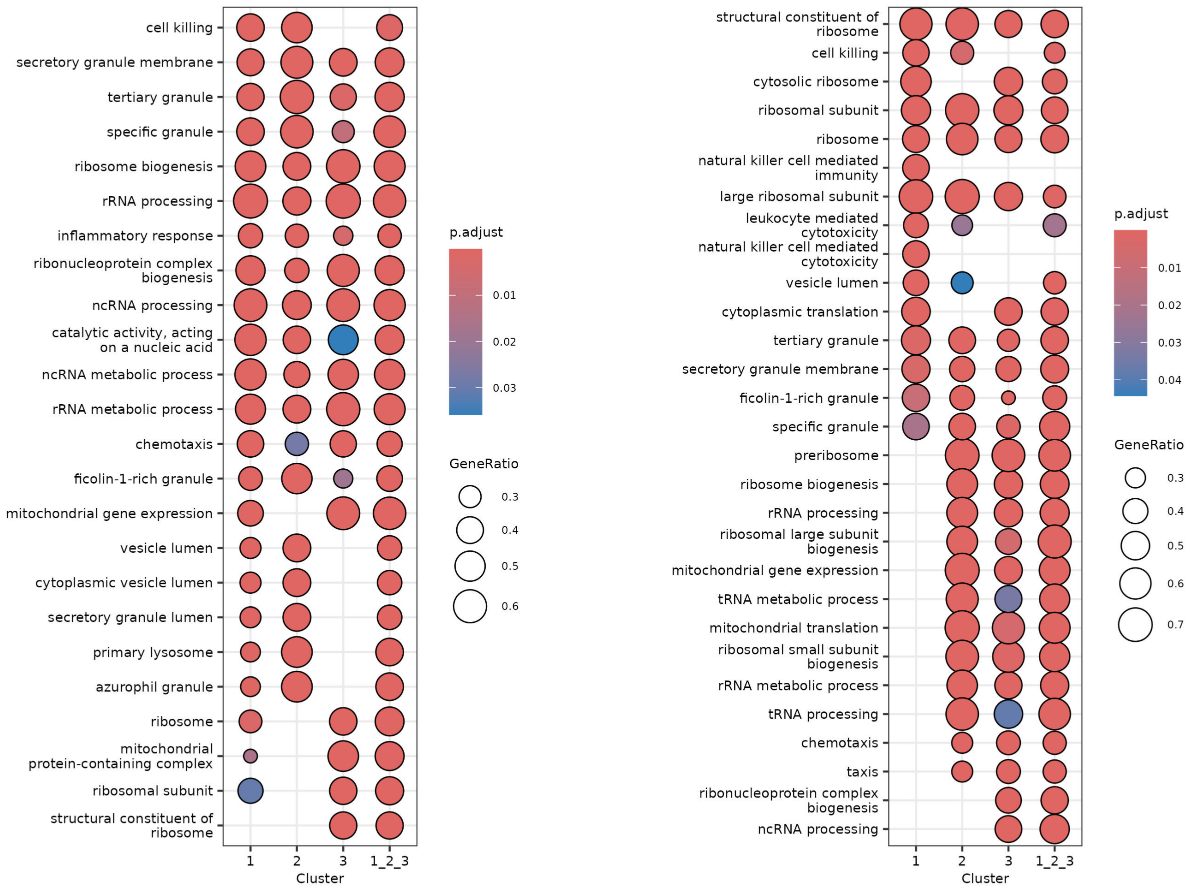


**Supplementary Figure 5**: Enrichment analysis of RNA molecules ranked using log2FC scores from Deseq2 analysis against HC for all clusters with differences in a) MDS-UPDRS3 and b) SFT. The most highly enriched pathways ranked by p-value are shown. The activated and suppressed pathways indicate a positive and negative enrichment score.

**Supplementary Table 1: Estimated proportions of cell types for the multi-omics clusters with differences in motor severity**
The table shows values for variables in year 3 (unless indicated as a future visit). The summary shows the median and inter-quartile range (IQR) in brackets. The cell type proportion were estimated from Whole Blood RNA sequencing data.

| | HC | Cluster 1 (Severe motor) | Cluster 2 (Intermediate motor) | Cluster 3 (Mild motor) | $p^a < 0.05$ |
|---|---|---|---|---|---|
| B.Memory | 2.88 | 1.92 | 2.88 | 1.92 | |
| B.Naive | 0.96 | 0.00 | 0.00 | 0.96 | * |
| Basophils.LD | 4.81 | 4.81 | 3.85 | 3.85 | * |
| MAIT | 2.88 | 2.88 | 2.88 | 2.88 | |
| mDCs | 0.00 | 0.00 | 0.00 | 0.00 | * |
| Monocytes.C | 14.42 | 12.50 | 14.42 | 13.46 | * |
| Monocytes.NC.I | 0.00 | 0.00 | 0.00 | 0.00 | |
| Neutrophils.LD | 37.50 | 40.38 | 42.31 | 42.31 | |
| NK | 0.96 | 0.96 | 0.96 | 0.96 | |
| pDCs | 0.00 | 0.00 | 0.00 | 0.00 | |
| T.CD4.Memory | 10.58 | 13.46 | 9.62 | 7.69 | * |
| T.CD4.Naive | 9.62 | 5.77 | 6.73 | 11.54 | * |
| T.CD8.Memory | 0.00 | 0.00 | 0.00 | 0.00 | |
| T.CD8.Naive | 9.62 | 9.62 | 8.65 | 8.65 | |
| T.gd.non.Vd2 | 0.00 | 0.00 | 0.00 | 0.00 | * |

[a]P-values were calculated using Kruskal-Wallis test (for continuous variables) and $\chi^2$ test (for categorical) between the three PD clusters.

**Supplementary Table 2: Estimated proportions of cell types for the multi-omics clusters with differences in cognitive severity**
The table shows values for variables in year 3 (unless indicated as a future visit). The summary shows the median and inter-quartile range (IQR) in brackets. The cell type proportion were estimated from Whole Blood RNA sequencing data.

| | HC | Cluster 1 (Intermediate cognitive) | Cluster 2 (Severe cognitive) | Cluster 3 (Mild cognitive) | $p^a < 0.05$ |
|---|---|---|---|---|---|
| B.Memory | 2.88 | 2.88 | 2.88 | 1.92 | * |
| B.Naive | 0.96 | 0.00 | 0.48 | 0.96 | * |
| Basophils.LD | 4.81 | 4.81 | 3.85 | 3.85 | * |
| MAIT | 2.88 | 3.85 | 2.88 | 2.88 | * |
| mDCs | 0.00 | 0.00 | 0.00 | 0.00 | * |
| Monocytes.C | 14.42 | 14.42 | 13.46 | 13.46 | |
| Monocytes.NC.I | 0.00 | 0.00 | 0.00 | 0.00 | |
| Neutrophils.LD | 37.50 | 39.42 | 43.27 | 42.31 | |
| NK | 0.96 | 0.96 | 0.00 | 0.96 | * |
| pDCs | 0.00 | 0.00 | 0.00 | 0.00 | |
| T.CD4.Memory | 10.58 | 13.46 | 9.13 | 7.69 | * |
| T.CD4.Naive | 9.62 | 4.81 | 7.69 | 11.54 | * |
| T.CD8.Memory | 0.00 | 0.00 | 0.00 | 0.00 | * |
| T.CD8.Naive | 9.62 | 9.62 | 9.13 | 8.17 | * |

[a]P-values were calculated using Kruskal-Wallis test (for continuous variables) and $\chi^2$ test (for categorical) between the three PD clusters.
